# Crystal structure of β-L-arabinobiosidase belonging to glycoside hydrolase family 121

**DOI:** 10.1101/2020.03.26.009860

**Authors:** Keita Saito, Alexander Holm Viborg, Shiho Sakamoto, Takatoshi Arakawa, Chihaya Yamada, Kiyotaka Fujita, Shinya Fushinobu

## Abstract

Enzymes acting on α-L-arabinofuranosides have been extensively studied; however, the structures and functions of β-L-arabinofuranosidases are not fully understood. Three enzymes and an ABC transporter in a gene cluster of *Bifidobacterium longum* JCM 1217 constitute a degradation and import system of β-L-arabinooligosaccharides on plant hydroxyproline-rich glycoproteins. An extracellular β-L-arabinobiosidase (HypBA2) belonging to the glycoside hydrolase (GH) family 121 plays a key role in the degradation pathway by releasing β-1,2-linked arabinofuranose disaccharide (β-Ara_2_) for the specific sugar importer. Here, we present the crystal structure of the catalytic region of HypBA2 as the first three-dimensional structure of GH121 at 1.85 Å resolution. The HypBA2 structure consists of a central catalytic (α/α)_6_ barrel domain and two flanking (N- and C-terminal) β-sandwich domains. A pocket in the catalytic domain appears to be suitable for accommodating the β-Ara_2_ disaccharide; this pocket is highly conserved among GH121 proteins. The three acidic residues Glu383, Asp515, and Glu713, located in this pocket, are completely conserved among all ~270 members of GH121; site-directed mutagenesis analysis showed that they are essential for catalytic activity. The active site of HypBA2 was compared with those of GH63 α-glycosidase, GH94 chitobiose phosphorylase, GH142 β-L-arabinofuranosidase, GH78 α-L-rhamnosidase, and GH37 α,α-trehalase. Based on these analyses, we concluded that the three conserved residues are essential for catalysis and substrate binding. β-L-Arabinobiosidase genes in GH121 are mainly found in the genomes of bifidobacteria and *Xanthomonas* species, suggesting that the cleavage and specific import system for the β-Ara_2_ disaccharide on plant hydroxyproline-rich glycoproteins are shared in animal gut symbionts and plant pathogens.

## Introduction

Enzymes and metabolic pathways involved in the microbial degradation of α-linked L-arabinofuranosyl (L-Ara*f*) residues, which are abundantly present in hemicellulosic polysaccharides of plants such as arabinoxylans and arabinans, have been extensively studied [1]. In the carbohydrate-active enzyme database [2], glycoside hydrolases (GHs) cleaving the α-L-Ara*f* bonds, including α-L-arabinofuranosidases (EC 3.2.1.55) and endo-1,5-α-L-arabinanases (EC 3.2.1.99), are classified into GH families 2, 3, 43, 51, 54, and 62. On the contrary, studies on enzymes cleaving β-linked L-Ara*f*s are limited, although such carbohydrate linkages have been found in hydroxyproline-rich glycoproteins (HRGPs) and peptide hormones [3–5]. Enzymes cleaving β-L-Ara*f* bonds were found in human symbiotic gut bacteria after 2011; these enzymes are associated with carbohydrate exploitation in the human gastrointestinal tract. At first, two enzymes were discovered from a human gut bacterium, *Bifidobacterium longum*, and classified into GH121 and GH127 [6,7]. Subsequent studies on one of the major human gut bacteria, *Bacteroides thetaiotaomicron*, led to of the discovery of other enzymes cleaving the β-L-Ara*f* bonds in plant pectins; these enzymes were classified as GH137, GH142, and GH146 [8,9]. The molecular mechanism of β-L-Ar*f* bond hydrolysis was solely studied on a GH127 β-L-arabinofuranosidase HypBA1 [10]. Although three-dimensional structures were determined for GH137, GH142, and GH146, the molecular mechanism of these enzymes was not uncovered in detail in the available literature [8,9].

*Bifidobacterium* species play a significant role in human health by acting with various transporters, glycosidases, and metabolic enzymes [11]. A gene cluster in the genome of *Bifidobacterium longum* JCM 1217 encodes three enzymes (Fig. 1) and a transporter involved in the degradation system for plant HRGPs [6,7]. The three enzymes belong to different GH families: GH43 α-L-arabinofuranosidase (HypAA, locus tag = BLLJ_0213), GH121 β-L-arabinobiosidase (HypBA2, BLLJ_0212), and GH127 β-L-arabinofuranosidase (HypBA1, BLLJ_0211). The transporter is an ABC-type sugar importer system that consists a substrate-binding protein (SBP, BLLJ_0208) and two transmembrane domain proteins (BLLJ_0209 and BLLJ_0210). Hydroxyproline (Hyp) residues in HRGPs, such as extensin and solanaceous lectins, are usually *O*-glycosylated with 3 or 4 L-Ara*f* residues that are linked with three β-bonds and one α-bond (Ara_3_-Hyp and Ara_4_-Hyp, Fig. 1) [12–14]. The chemical structures of Ara_3_-Hyp and Ara_4_-Hyp are Ara*f*-β1,2-Ara*f*-β1,2-Ara*f*-β-Hyp and Ara*f*-α1,3-Ara*f*-β1,2-Ara*f*-β1,2-Ara*f*-β-Hyp, respectively. The three enzymes in the bifidobacterial β-L-arabinooligosaccharide degradation pathway consist of two extracellular enzymes and one intracellular enzyme. The two extracellular enzymes (HypAA and HypBA2) initially act on the β-L-arabinooligosaccharides. HypAA releases the terminal L-arabinose (L-Ara) from Ara_4_-Hyp by cleaving the α1,3-bond (unpublished data), and HypBA2 liberates Ara*f*-β1,2-Ara*f* (β-Ara_2_) from Ara_3_-Hyp [6]. The ABC transporter likely imports β-Ara_2_ because the SBP (BLLJ_0208) specifically binds β-Ara_2_ (Miyake et al., paper in revision). An intracellular enzyme, HypBA1, cleaves β-Ara_2_ to release two L-Ara molecules [7].

**Fig. 1.**
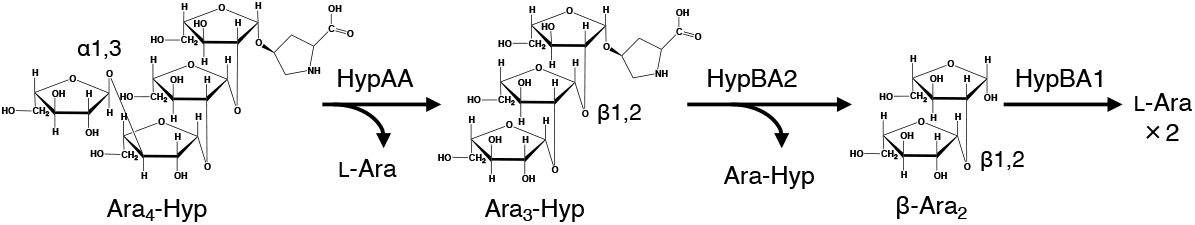
Glycosidic bond cleavage of the β-arabinooligosaccharide degradation system of *B. longum* JCM 1217. HypAA and HypBA2 are extracellular enzymes, and HypBA1 is an intracellular enzyme. An ABC-type transporter system specifically imports β-Ara_2_ into the cell.

The GH121 β-L-arabinobiosidase HypBA2 plays a key role in the degradation system by releasing the β-Ara_2_ that is transferred to the specific sugar importer. GH121 currently consists of 281 bacterial ORFs (as of March 2020), and HypBA2 is the sole characterized member. In this study, we report the crystal structure of HypBA2 as the first three-dimensional structure of GH121.

## Materials and Methods

### Protein expression and purification

The expression plasmid for C-terminally His-tagged proteins of CΔ789 (residues 33–1154) and CΔ1049 (residues 33–894) were constructed in our previous study [6]. Selenomethionine (SeMet)-labeled and native proteins were expressed in *Escherichia coli* BL21-CodonPlus (DE3)-RP-X and BL21-CodonPlus (DE3)-RIL (Agilent Technologies, Santa Clara, CA, USA), respectively. The transformants were grown at 37 °C for 5 h in LeMaster medium (SeMet-labeled protein) or Luria-Bertani medium (native protein) containing 50 μg/mL ampicillin and 34 μg/mL chloramphenicol. Protein expression was induced by adding 0.1 mM isopropyl-β-D-thiogalactopyranoside to the medium, and the cells were further cultivated at 25 °C for 20 h. The cells were harvested by centrifugation and suspended in 20 mM Tris-HCl (pH 8.0), 250 mM NaCl, and 2 mM CaCl_2_ (buffer A). The cells were disrupted via sonication, and the supernatant was purified by sequential column chromatography. Ni-affinity chromatography was conducted using a HisTrap FF column (GE Healthcare, Fairfield, CT, USA) with wash and elution steps of 20 mM and 400 mM imidazole in Buffer A, respectively. The eluted protein sample was dialyzed against 20 mM Tris-HCl (pH 8.0) and 2 mM CaCl_2_ (buffer B) and applied to a Mono Q 10/100 GL column (GE Healthcare) equilibrated with buffer B. The protein sample was eluted with a linear gradient of 0 to 1 M NaCl. The protein sample was concentrated by ultrafiltration (Amicon Ultra-4 centrifugal filter devices, 50,000 MWCO; Millipore, Billerica, MA, USA), and the solution was changed to buffer A. Gel filtration chromatography was conducted using a HiLoad 16/60 Superdex 200 pg column (GE Healthcare) equilibrated with buffer A at a flow rate of 1 mL/min. The purified protein was again concentrated by ultrafiltration, and the solution was changed to a buffer consisting of 5 mM Tris-HCl (pH 8.0) and 2 mM CaCl_2_. Protein concentrations were determined by a BCA protein assay kit (Thermo Fisher Scientific, Waltham, MA, USA) with bovine serum albumin as the standard.

### Crystallography and visualization

The protein crystals were grown at 20 °C using the sitting drop vapor-diffusion method by mixing 0.5 μL of protein solution with an equal volume of a reservoir solution. The SeMet-labeled CΔ789 was crystallized using a protein solution (12.5 mg/mL) and a reservoir solution containing 17% (w/v) PEG1000 and 0.1 M Tris-HCl (pH 6.6). The native CΔ1049 protein was crystallized using a protein solution (5 mg/mL) and a reservoir solution containing 40% (w/v) PEG400 and 0.1 M Na-acetate (pH 4.5). Crystals were flash-cooled by dipping into liquid nitrogen. X-ray diffraction data were collected at 100 K on beamlines at the Photon Factory of the High Energy Accelerator Research Organization (KEK, Tsukuba, Japan). Preliminary diffraction data were collected at SPring-8 (Hyogo, Japan). The data sets were processed using XDS [15]. The phase determination and automated model building were performed using PHENIX [16]. Manual model rebuilding and crystallographic refinement were performed using Coot [17] and Refmac5 [18]. Molecular graphic images were prepared using MyMOL (Schrödinger, LLC, New York, NY, USA). A structural similarity search was performed using the Dali server (http://ekhidna2.biocenter.helsinki.fi/dali/) [19]. Sequence conservation mapping was performed using the ConSurf server [20].

### Construction of site-directed mutants and enzyme assay

Site-directed mutants were constructed using PrimeSTAR Max DNA Polymerase (Takara Bio Inc., Shiga, Japan). The following primers and their complementary primers were used: 5’- TCCATCgcgGGTGTGCTCGGCTACAAC-3’ for E383A, 5’-TCCATCcagGGTGTGCTCGGCTACAAC-3’ for E383Q, 5’-GGTAACgcgGCCGACGCCGTTTCCTTC-3’ for D515A, 5’-GGTAACaacGCCGACGCCGTTTCCTTC-3’ for D515N, 5’-CAAAACgcgTTCTGGAACGAGGACAAC-3’ for E713A, and 5’-CAAAACcagTTCTGGAACGAGGACAAC-3’ for E713Q (mutated sites shown with lowercase letters).

The activity assay was performed using *cis*-Ara_3_-Hyp-DNS as the substrate. A reaction mixture containing 50 μM Ara_3_-Hyp-DNS, 5 mM Na-acetate buffer (pH 5.5), and HypBA2 protein was incubated at 37 °C for 1 h. Two μM of the reaction mixture was spotted onto a thin layer chromatography (TLC) Silica Gel 60 F_254_ plate (Merck, Darmstadt, Germany). Subsequently, the TLC plate was developed by a 2:1:1 solvent mixture (v/v/v) of ethyl acetate/acetic acid/water. Fluorescence of the dansyl (DNS) group was detected under a UV lamp (365 nm).

### Sequence analysis and molecular phylogeny

269 GH121 sequences were extracted from the CAZy database (January 6^th^, 2020) and preliminarily aligned using MAFFT [21]. The multiple sequence alignment was inspected in Jalview [22], and all sequences were cut according to the observed boundaries of the conserved GH121 module. These modules were then realigned with MAFFT G-INS-i (iterative refinement using pairwise Needleman–Wunsch global alignments). A maximum likelihood phylogenetic tree was estimated with RAxML [23] using 100 bootstrap replicates and visualized with iTOL [24]. Short and high bootstrap-supported branches in the phylogenetic tree were collapsed, and major features and number of sequences were displayed.

## Results

### Identification and structure determination of the catalytic domain

HypBA2 is a multi-domain protein with 1943 amino acids (aa) (Fig. 2A). Presence of the N-terminal signal sequence and the C-terminal transmembrane region indicate that this enzyme is anchored to outside of the cell of the gram-positive bacterium *(B. longum)*. In our previous study, the N-terminal half of BLLJ_0212 was identified as the conserved region of GH121, and a deletion analysis indicated that a region of residues 33–1051 showed full activity in the presence of 1 mM Ca^2+^ [6]. Protein purification of HypBA2 deletants was conducted in the presence of 2 mM CaCl_2_ throughout the steps (see Materials and Methods). We tried to crystallize various deletion constructs of HypBA2 and succeeded in solving the crystallographic structure using a SeMet-derivative of a construct of residues 33–1154 (CΔ789) by the single-wavelength anomalous dispersion method (Fig. 2A and Table 1). Automated model building and subsequent manual model building resulted in a SeMet-containing protein model with many disordered regions, and we could not build a reliable polypeptide model in the C-terminal region (residues 896–1154) due to ambiguous electron density (data not shown). In our previous study, a construct of residues 33–894 (CΔ1049, see Fig. 2A) showed about 16% activity compared to the full-length enzyme [6]. Our re-analysis using the purified samples of the CΔ789 and CΔ1049 deletants demonstrated that both of them exhibited the catalytic activity of cleaving Ara_3_-Hyp-DNS to release Ara-Hyp-DNS (Fig. 3, left). Therefore, we also crystallized the native protein of CΔ1049. The purified protein samples of CΔ789 and CΔ1049 migrated as a single band on SDS-PAGE that was consistent with their calculated molecular masses (124,555 and 96,854 Da, respectively), and gel filtration experiments suggested that the recombinant proteins are both monomeric in solution (data not shown). The crystal structure of the native (unlabeled) CΔ1049 protein was determined at 1.85 Å resolution (Table 1). The refined crystal structure of CΔ1049, which contains the core catalytic region of GH121, comprises residues 38–894, and three short regions (residues 438–445, 512–515, and 889–891) were disordered (Fig. 2A). Two residues from a C-terminal His_6_-tag (LE of LEHHHHHH) were visible in the crystal structure and modeled as residues 895 and 896. Extensive attempts at co-crystallization and soaking experiments with β-Ara_2_ (the reaction product) were undertaken, but no electron densities were found in the putative active site (data not shown).

**Fig. 2.**
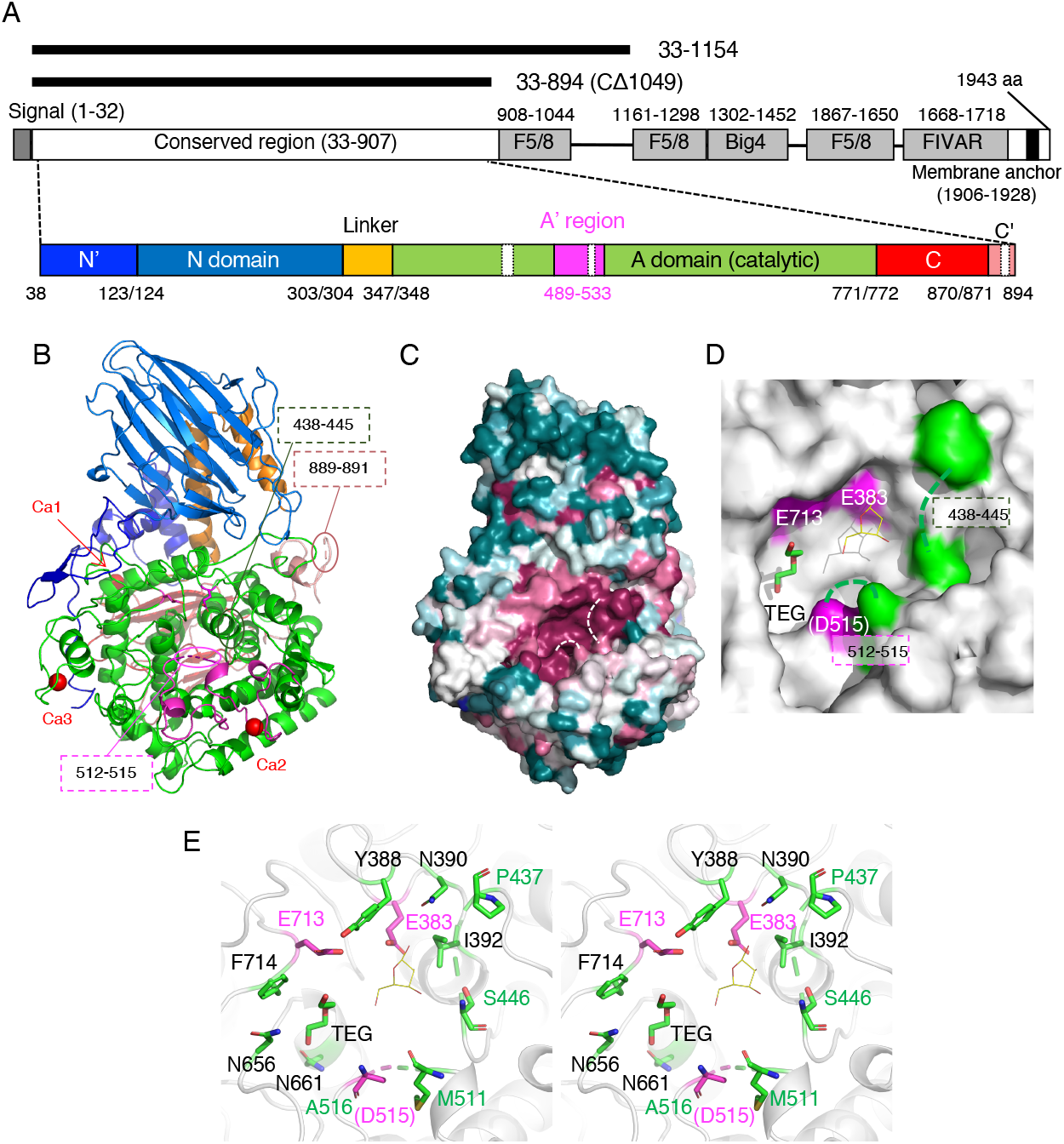
Structural architecture of HypBA2. (A) Schematic representations of the full-length domain structure of HypBA2 are shown in the middle. Regions and domains modeled in the crystal structure are schematically shown below. Deletion constructs used for crystallography are shown above. (B) The overall structure of the CΔ1049 construct (residues 33–894) of HypBA2. (C) Sequence conservation mapping on the molecular surface. Amino acid sequence conservation among GH121 is colored with red (high), white (middle), and blue (low). (D) Molecular surface representation of the active site pocket in the catalytic domain. Glu383, Asp515, and Glu713 are conserved putative catalytic residues of GH121 (shown in magenta). Because Asp515 is disordered, the neighboring residue (Asp516) is shown. A triethylene glycol molecule (TEG) bound to HypBA2 and a β-L-Ara*f* molecule bound to GH142 BT_1020 are shown as green sticks and thin yellow lines, respectively. (E) Stereoview of the active site of HypBA2. (A–E) Disordered regions are indicated with dotted lines.

**Fig. 3.**
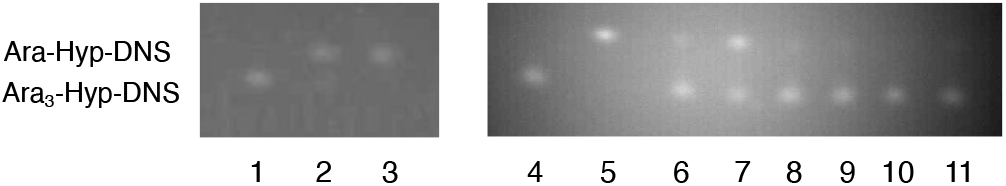
TLC analysis of the activities of deletion constructs and site-directed mutants towards Ara_3_-Hyp-DNS. The reaction was performed in a solution containing 50 μM Ara_3_-Hyp-DNS (substrate) and 50 mM Na-acetate buffer (pH 5.5). After development, the fluorescence of the dansyl group was detected. (Left) The substrate was incubated either without *(lane 1)* or with the deletion constructs *(lanes 2 and 3*, 0.45 mg/ml protein) at 30 °C for 20 min. *Lane 2*, CΔ1049; *lane 3*, CΔ789. (Right) The substrate was incubated either without *(lane 4)*, with CΔ1049 *(lane 5*, 2.5 mg/ml protein), or with site-directed point mutants of CΔ1049 *(lanes 6-11*, 2.5 mg/ml protein) at 37 °C for 1 h. *Lane 6*, D515A; *lane 7*, D515N; *lane 8*, E713A; *lane 9*, E713Q; *lane 10*, E383A; *lane 11*, E383Q.

**Table 1.** Data collection and refinement statistics of the crystallography of HypBA2.

| Data set | SeMet protein (33-1154) | Native protein (33-894) |
| --- | --- | --- |
| Data collection <sup>a</sup> |  |  |
| Beamline | NW12A | BL1A |
| Wavelength (Å) | 0.9791 | 1.1000 |
| Space group | $P2_12_12_1$ | $P2_12_12_1$ |
| Unit cell (Å) | $a = 76.10, b = 100.60, c = 141.16$ | $a = 81.53, b = 88.62, c = 127.92$ |
| Resolution (Å) | 50.0–2.20<br>(2.24–2.20) | 44.31–1.85<br>(1.89–1.85) |
| Total reflections | 1,641,170 | 542,053 |
| Unique reflections | 106,306 (5,342) | 79,741 (4,473) |
| Completeness (%) | 100.0 (100.0) | 100.0 (100.0) |
| Multiplicity | 29.4 (29.0) | 6.8 (7.0) |
| Mean $I/\sigma(I)$<sup>b</sup> | 28.6 (4.4) | 9.1 (1.3) |
| $R_{\text{merge}}$ | 0.119 (0.667) | 0.123 (1.405) |
| $CC_{1/2}$ | (0.928) | 0.940 (0.634) |
| Anomalous completeness (%) | 100.0 (100.0) |  |
| Anomalous multiplicity | 15.4 (15.0) |  |
| Refinement |  |  |
| Resolution (Å) |  | 44.35–1.85 |
| No. of reflections |  | 75,729 |
| $R_{\text{work}}/R_{\text{free}}$ | | 0.158/0.185 |
| Number of atoms |  | 7,337 |
| RMSD from ideal values |  |  |
| Bond lengths (Å) |  | 0.011 |
| Bond angles (°) |  | 1.633 |
| Ramachandran plot (%) |  |  |
| Favored/allowed/outlier |  | 97.4/2.5/0.1 |
| PDB code |  | 6M5A |

<sup>a</sup> Values in parentheses are for the highest resolution shell.

### Crystal structure of GH121 HypBA2

The overall structure of CΔ1049 consists of a mostly unstructured N-terminal region (residues 38–123, blue), a N-domain (124–303, marine), a linker region with two α-helices (304–347, orange), a catalytic (α/α)_6_ barrel domain (348–771, green), a C-domain (772–870, red), and a mostly unstructured C-terminal region with an α-helix (871–894, pink; Fig. 2B). The N- and C-domains adopt a β-sandwich fold. The N-domain is similar to an existing CATH domain (Superfamily 2.70.98.50 with Dali Z score ~ 11) [25] while the C-domain shows no similarity to established protein folds (Dali Z-score < 4). A Dali structural similarity search analysis of the whole structure showed that the catalytic region of GH121 is primarily similar to the GH142 protein (BT_1020, Table 2). Subsequent hits were GH enzymes belonging to GH63, GH78, GH94, and GH37, and all of them adopt a catalytic (α/α)_6_ barrel domain (discussed below). Interestingly, all of the structurally similar GH families are inverting enzymes, whereas GH121 HypBA2 is a retaining enzyme because it exhibited transglycosylation activity [6]. The catalytic barrel region of a structural homolog in GH63 (α-glycosidase YgjK from *E. coli*) is designated as the “A-domain,” and an extra structural unit within this domain is designated as the “A’-region” [26]. The A’-region is a long insertion between the fifth and sixth helices in the twelve helices of the (α/α)_6_ barrel scaffold. In HypBA2, a similar insertion corresponding to the A’-region is also present (magenta in Fig. 2A and 2B, residues 489–533).

**Table 2.**
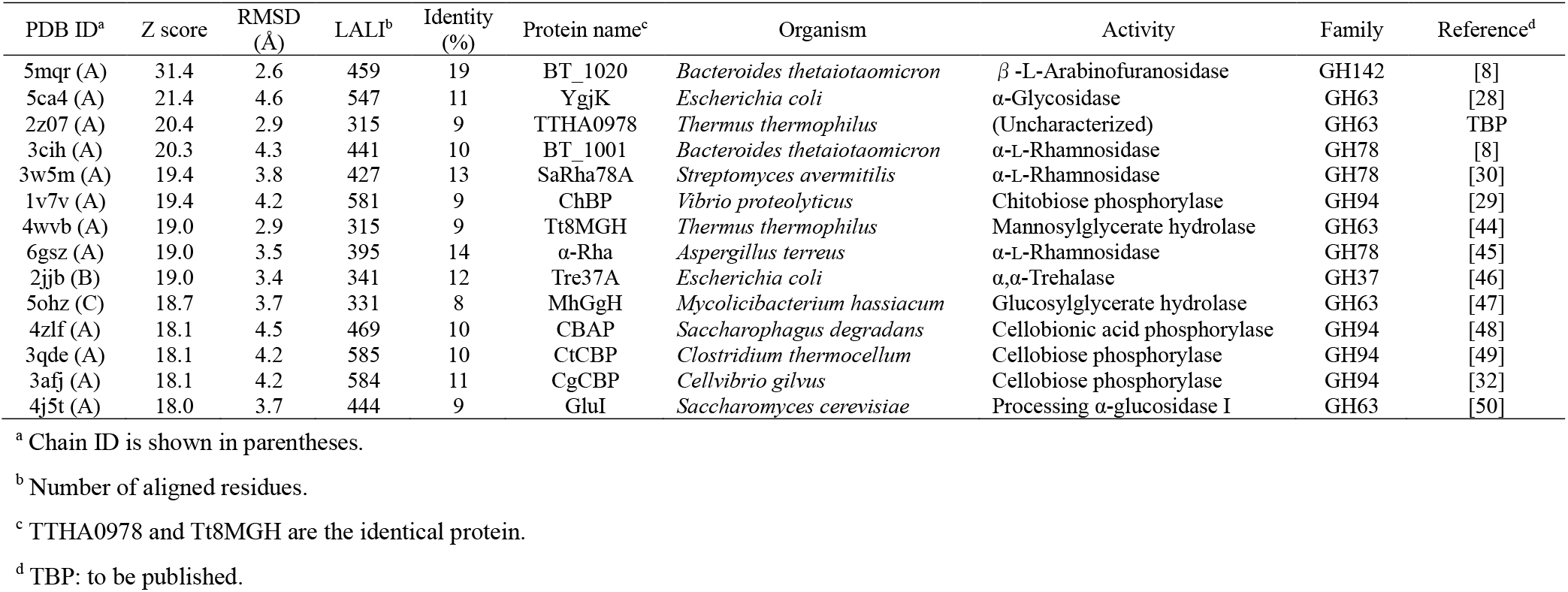
Result of structural similarity search using the Dali server.

The crystal structure of CΔ1049 contains three calcium ions (Fig. 2B, red spheres). The first Ca^2+^ (Ca1) is located between the linker region and the catalytic domain and coordinated by the side chains of Asp345, Asp349, and Asp352, the main chain carbonyl of Thr346, and two water molecules. The second Ca^2+^ (Ca2) is located in the A’-region and coordinated by the side chains of Asn497, Asn499, Asn501, Asp505, and Asp532, and the main chain carbonyl of Leu503. The third Ca^2+^ (Ca3) is located in a C-terminal region of the catalytic domain and coordinated by the side chains of Asp724, Asn726, Asp728, and Asp735, the main chain carbonyl of Val730, and one water molecule.

### Active site structure and site-directed mutant analysis

The degree of amino acid sequence conservation of GH121 members mapped on the molecular surface of HypBA2 clearly illustrated a highly conserved pocket on the catalytic domain (Fig. 2C, red). Three conserved acidic residues (Glu383, Asp515, and Glu713) of GH121 (discussed below) and two short disordered regions (438–445 and 512–515) are located in the pocket (Fig. 2D). Because one of the three conserved acidic residues (Asp515) was disordered in the crystal structure, the neighboring residue (Ala516) is shown with a magenta color in the figures. A triethylene glycol (TEG) molecule, which was derived from the crystallization precipitant, was bound in the pocket (a green stick in Fig. 2D). A superimposed β-L-Ara*f* molecule bound to GH142 BT_1020 is shown as thin yellow lines (discussed below), illustrating a possible subsite –1 area in the pocket. Glu383 and Glu713 are located near the superimposed β-L-Ara*f* molecule (Fig. 2E). Asp516 is located in a relatively far position, but Asp515 can approach the catalytic area with the flexibility of the disordered region.

Then, we constructed point mutants of CΔ1049 by site-directed mutagenesis. As shown in Fig. 3 (right), mutations of Glu383 and Glu713 (E383A, E383Q, E713A, and E317Q) completely abolished the activity. On the other hand, mutants of Asp515 showed very weak activity, and substitution with Asn (D515N) appeared to exhibit higher activity than an Ala mutant (D515A).

### Sequence analysis of GH121

We conducted a protein sequence analysis of the 272 GH121 entries listed on the CAZy database. After the deletion of 3 duplicates, 269 entries were analyzed by MAFFT multiple sequence alignment, and 65 of them are shown in Supplementary Fig. S1. A phylogenetic tree (cladogram) indicated that HypBA2 is in a cluster of 22 bifidobacterial sequences (Fig. 4A). 186 sequences from *Xanthomonas* species formed a very large group with little or no sequence diversity, 175 of them being in a zero branch between them. A HypBA2 homolog from *Xanthomonas euvesicatoria* (XeHypBA2) [27] was also included in this cluster. Sequence logos around the three acidic residues (E383, D515, and E713, Fig. 4B) illustrated that all of them are completely conserved in GH121, and nearby sequences are also highly conserved. Therefore, the structural feature of the active site pocket, the site-directed mutagenesis analysis, and the comprehensive sequence analysis strongly suggested that the three acidic residues play important roles in the catalytic process and/or substrate binding of HypBA2 and GH121 enzymes.

**Fig. 4.**
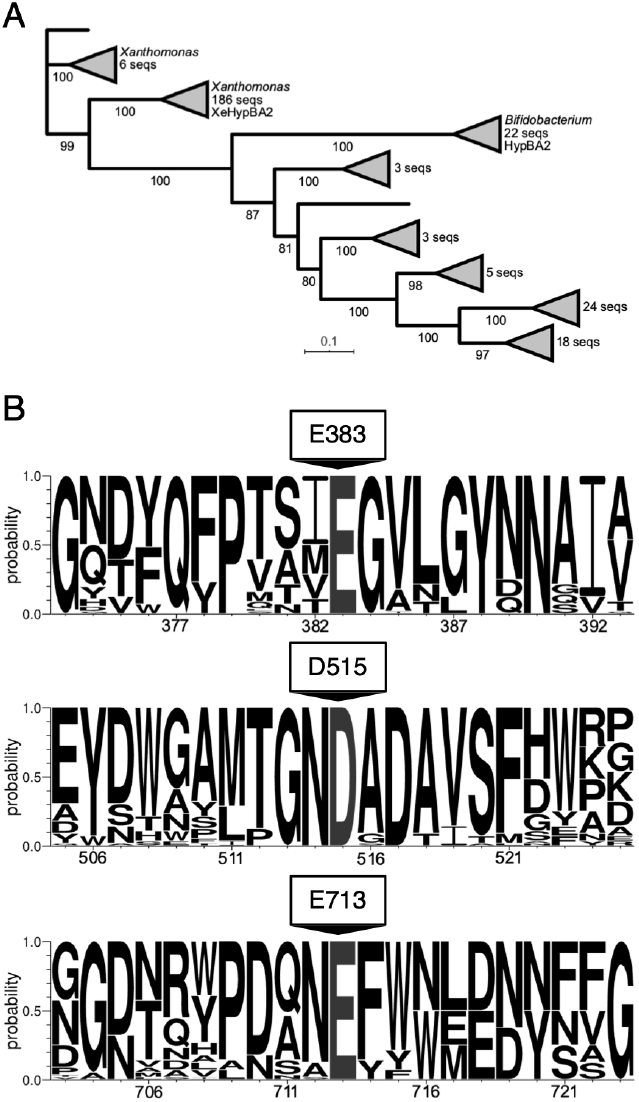
A phylogenetic tree (A) and sequence conservation around the selected residues (B) of GH121. (A) The maximum likelihood phylogenetic tree was generated using 269 GH121 catalytic domain sequences. Bootstrap values based on 100 replicates are shown. Short and high bootstrap support branches are collapsed as triangles. (B) Sequence logo around Glu383, Asp515, and Glu713 indicated that these residues are completely conserved in GH121.

### Structural comparison with other GH families

Then, we examined possible catalytic residues of GH121 by comparing its structure with structural homologs. The overall structure of HypBA2 (Fig. 5A) was compared with representative structures of GH63 (Fig. 5B, α-glycosidase YgjK from *E. coli)* [28], GH94 (Fig. 5C, chitobiose phosphorylase ChBP from *Vibrio proteolyticus)* [29], GH142 (Fig. 5D, β-L-arabinofuranosidase BT_1020 from *B. thetaiotaomicron*) [8], GH78 (Fig. 5E, α-L-rhamnosidase SaRha78A from *Streptomyces avermitilis)* [30], and GH37 (Fig. 5F, α,α-trehalase Tre37A from *E. coli*) [31]. The GH63 and GH94 enzymes share multi-domain structures containing the N-domain (marine in Fig. 5), the linker region (orange), and the catalytic barrel domain (green). GH94, GH14, and GH78 have a β-sandwich domain similar to the C-domain (red) behind the barrel domain. It is noteworthy that GH63 has a calcium ion (Fig. 5B, red sphere) at a similar position to Ca2 of HypBA2. GH37 only has a catalytic barrel domain as a structurally conserved element. The catalytic general base and acid residues for the anomer-inverting glycosidic bond hydrolysis in GH63, GH78, and GH37 have been identified in previous reports [28,30,31]. The general base residue of all of these families is located at the position corresponding to Glu713 in HypBA2 (Fig. 5A, B, E, and F). The general acid residue of GH63 and GH37 is located in the A’-region (magenta), corresponding to the position of Asp515 in HypBA2, while the general acid residue of GH78 appears to be located at a position corresponding to Glu383 in HypBA2. GH94 enzymes are inverting phosphorylases that use an inorganic phosphate as a nucleophile [29,32]. A sulfate molecule (phosphate analog) and the general acid residue of GH94 are located at positions corresponding to Glu713 and Asp515 in HypBA2, respectively (Fig. 5A and C). The catalytic residue of GH142 was not identified, but a conserved residue (Glu694) near the β-L-Ara*f* molecule in the putative catalytic pocket was identified (Fig. 5D) [29].

**Fig. 5.**
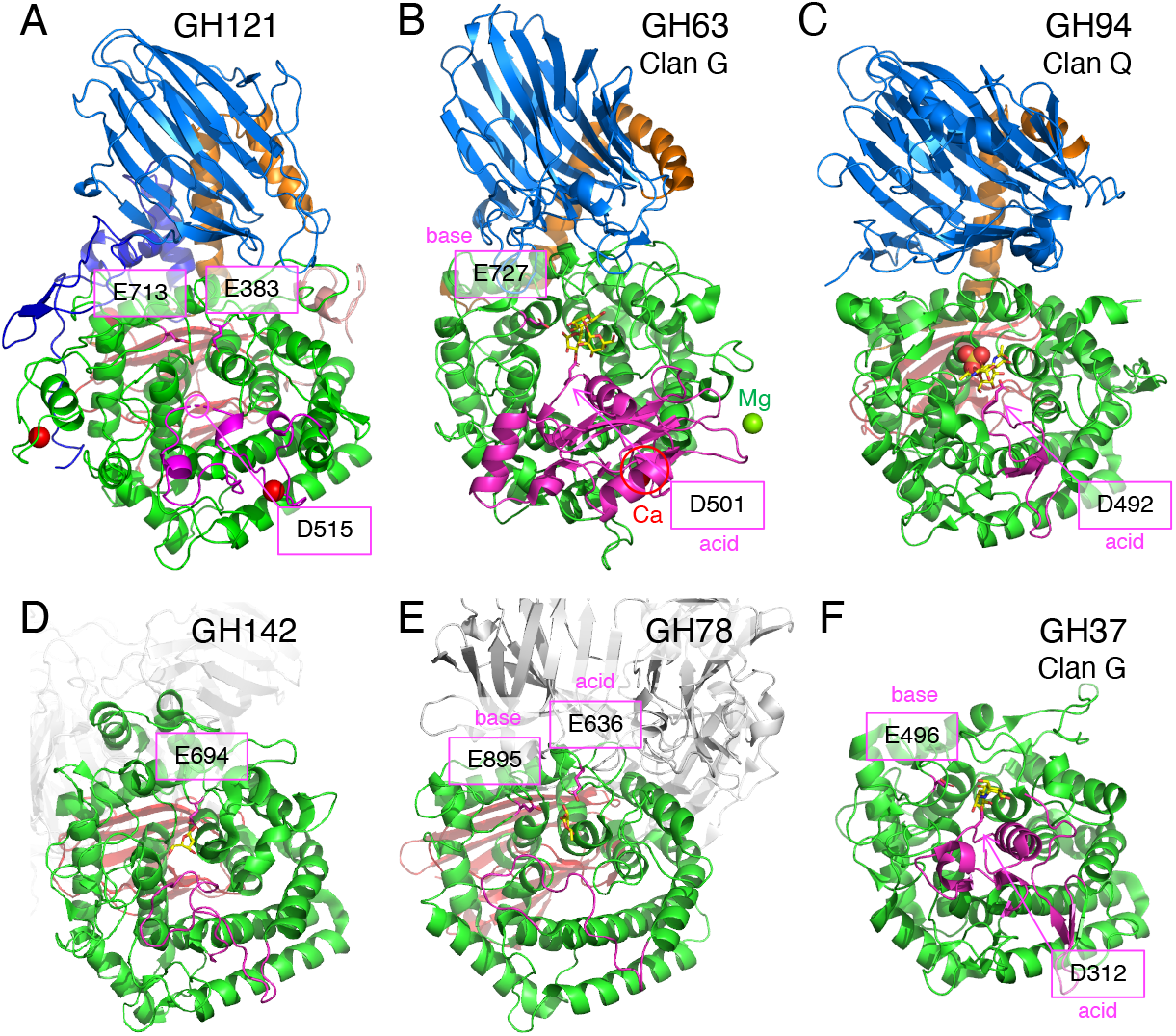
Structural comparison of HypBA2 with structural homologs belonging to other GH families. Overall structures of (A) GH121 HypBA2, (B) GH63 α-glycosidase YgjK from *E. coli* (PDB ID 5CA3), (C) GH94 chitobiose phosphorylase ChBP from *V. proteolyticus* (1V7X), (D) GH142 β-L-arabinofuranosidase BT_1020 from *B. thetaiotaomicron* (5MQS), (E) GH78 α-L-rhamnosidase SaRha78A from *S. avermitilis* (3W5N), and (F) GH37 α,α-trehalase Tre37A from *E. coli* (2JF4) are shown. Catalytic residues are indicated with a magenta color. The A’-region (insertion between the fifth and sixth helices of the barrel) is also shown with a magenta color.

Fig. 6A shows a superimposition of GH142 β-L-arabinofuranosidase BT_1020 (thick sticks) and HypBA2 (thin sticks) at the active site. Glu383 in HypBA2 is overlaid well with the putative catalytic residue of BT_1020 (Glu694, red sticks). Despite the ambiguous electron density and low crystallographic resolution (3.0 Å) [8], we consider that the position of the β-L-Ara*f* molecule indicates the approximate location of the subsite –1 in BT_1020. Glu694 forms bidentate hydrogen bonds with the O1 and O2 hydroxyls of the β-L-Ara*f* molecule. Tyr702, Gly805, Trp943, and His1019 are also located near the ligand, but they are not conserved in GH121 at all. As shown in Fig. 6B, the catalytic base and acid residues of GH63 enzyme (YgjK) correspond to Glu713 and Asp515 in HypBA2. The position of Asp501 (general acid) in YgjK is located above the subsite –1 sugar as an “*anti*” protonator according to the definition by Nerinckx et al. [33]. The catalytic base residue in GH78 (Glu895 in SaRha78A) also corresponds to Glu713 in HypBA2 (Fig. 6C). However, the location of the general acid residue of GH78 (Glu636) does not correctly match the position of Glu383 in HypBA2 but rather is located significantly “above” it. It is noteworthy that the general acid residues of GH63, GH78, and GH37 are all “*anti*” positioned (see the “syn/anti lateral protonation” page in CAZypedia, https://www.cazypedia.org) [34]. In comparison with GH142, GH63, and GH78 enzymes, Glu383 of GH121 appears to form direct interactions with the subsite –1 sugar. In GH13 enzymes, the anchor (or fixer) is the third conserved residue of this large and divergent family and plays a critical role in substrate binding by forming bidentate hydrogen bonds with the α-glucoside at subsite –1 [35–37]. The anchor residue ensures nucleophilic and acid/base catalysis on the glycosidic bond by proper positioning of the substrate.

**Fig. 6.**
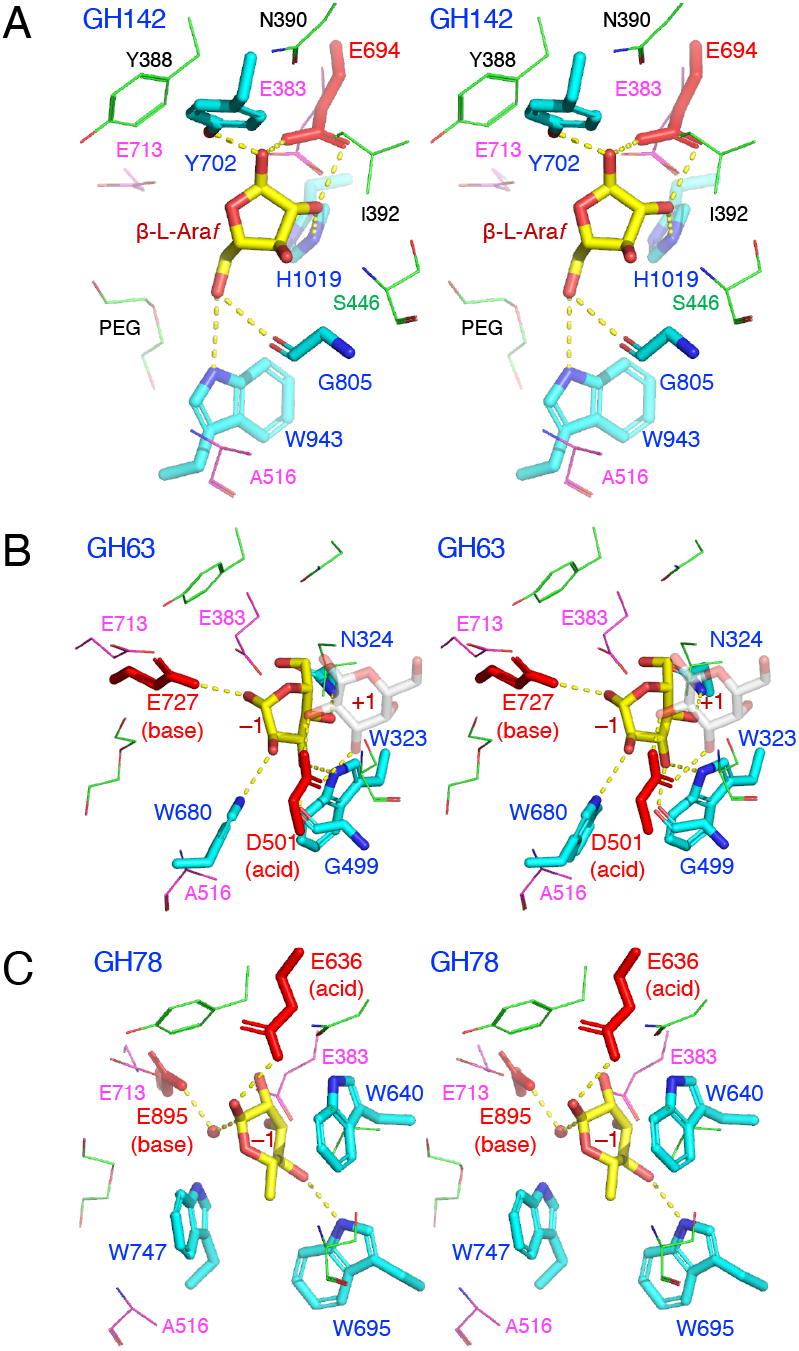
Stereoview of the active site of structural homologs belonging to other GH families. (A) GH142 β-L-arabinofuranosidase BT_1020 (cyan) complexed with β-L-Ara*f* (yellow), (B) GH63 *α*-glycosidase YgjK (cyan) complexed with glucose (yellow) and lactose (galactose at subsite +1 is shown as transparent grey sticks), and (C) GH78 α-L-rhamnosidase SaRha78A (cyan) complexed with α-L-rhamnose (yellow). The catalytic residues of BT_1020, YgjK, and SaRha78A are shown as red sticks. Residues in the active site of HypBA2 are superimposed as magenta (putative catalytic residues) or thin green lines.

## Discussion

Based on structural comparisons, we suggest that Glu713 and Asp515 are the likely catalytic nucleophile and acid/base residues of HypBA2, respectively. This hypothesis assumes a proper positioning of the putative acid/base residue (Asp515) on substrate binding. The catalytic acid/base residue of GH29 1,3-1,4-α-L-fucosidase AfcB from *B. longum* subsp. *infantis* suggestively exhibited a significant movement on the substrate binding due to the flexibility of a loop carrying that residue [38]. However, this hypothesis is solely based on the structural comparison with other inverting GH families and a simple activity assay of the site-directed mutants. Further analyses, such as determination of ligand complex structures and detailed kinetic measurements, will identify the catalytic residues.

HypBA2 is an exo-type glycoside hydrolase that releases a disaccharide (β-Ara_2_). This mode of action is similar to other extracellular glycosidases of bifidobacteria involved in the degradation of human milk oligosaccharides and mucin *O*-glycans, *vis*. GH20 and GH136 lacto-*N*-biosidases [39,40] and GH101 endo-α-*N*-acetylgalactosaminidase [41]. The molecular surface representation of HypBA2 delineates a pocket that can accommodate the β-Ara_2_ disaccharide (Fig. 2D). The TEG molecule in the pocket appears to indicate the location of subsite –2. The CASTp server [42] calculated that the surface-accessible volume and area of the pocket are 960 Å^3^ and 685 Å^2^, respectively. The pocket size may be overestimated due to the two disordered regions. Given the hypothesis that one of the catalytic residues (Asp515) is located in the shorter disordered region (residues 512–515), the longer disordered region (residues 438–445) may cover and cap the active site pocket on substrate binding as in the case of GH127 HypBA1 [10]. Interestingly, Pro437, Gly438, and Trp443 are completely conserved in GH121 members (Supplementary Figure S1), suggesting loop flexibility near the Gly residue and a sugar-aromatic stacking interaction by the Trp side chain with the substrate.

In this study, we presented the first three-dimensional view of a GH121 enzyme and demonstrated that it has some structural relevance with several GH families, including GH63 and GH142. Since we could not find the closest structural and functional relative of GH121 among the GHs, the molecular evolution of GH121 enzymes are still enigmatic. Interestingly, most of the ~270 GH121 members are found only in bifidobacteria (animal gut symbionts, 22 sequences) and *Xanthomonas* species (plant pathogens, ~190 sequences, Fig. 4A). In contrast, GH127 and GH146 β-L-arabinofuranosidases (monosaccharide-releasing exo-enzymes), which were classified together as DUF1680 in Pfam [43], are widely distributed among various bacteria [7]. Bifidobacteria and *Xanthomonas* species apparently have no biological relationships with distinct biological niches. Bifidobacteria are representative species of gut microbes that are beneficial to human health while *Xanthomonas* species contain pathogens that cause bacterial spots on a wide variety of plant species. The genes for β-L-arabinooligosaccharide-degradation enzymes in *Xanthomonas euvesicatoria (xehypBA1-BA2-AA)* are not involved in either pathogenicity or non-host resistance reactions [27]. However, the distribution of GH121 genes suggests that bifidobacteria and *Xanthomonas* species possibly benefit from the disaccharide-releasing enzyme and the disaccharide-specific importer system on their biological niches in order to live on special types of plant glycans, such as Ara_3_-Hyp β-L-arabinooligosaccharides on HRGPs.

## Acknowledgments

We thank Dr. Akihiro Ishiwata for the helpful discussions, Mr. Masayuki Miyake for the preliminary crystallization screenings, and the staff of SPring-8 and KEK-PF for the X-ray data collection.

## Author Contributions

Conceptualization, Project Administration, and Supervision: SF KF. Investigation: KS AHV SS TA. Data Curation: SF TA KF. Formal Analysis: AHV. Funding Acquisition: SF TA CY KF. Resources: TA CY. Validation: SF TA KF. Writing – Original Draft Preparation: SF AHV Writing – Review & Editing: all authors.

## Data Availability

All relevant data are within the paper and its Supporting Information files except for those submitted to PDB (ID code 6M5A).

---

**Supplementary Figure S1.**
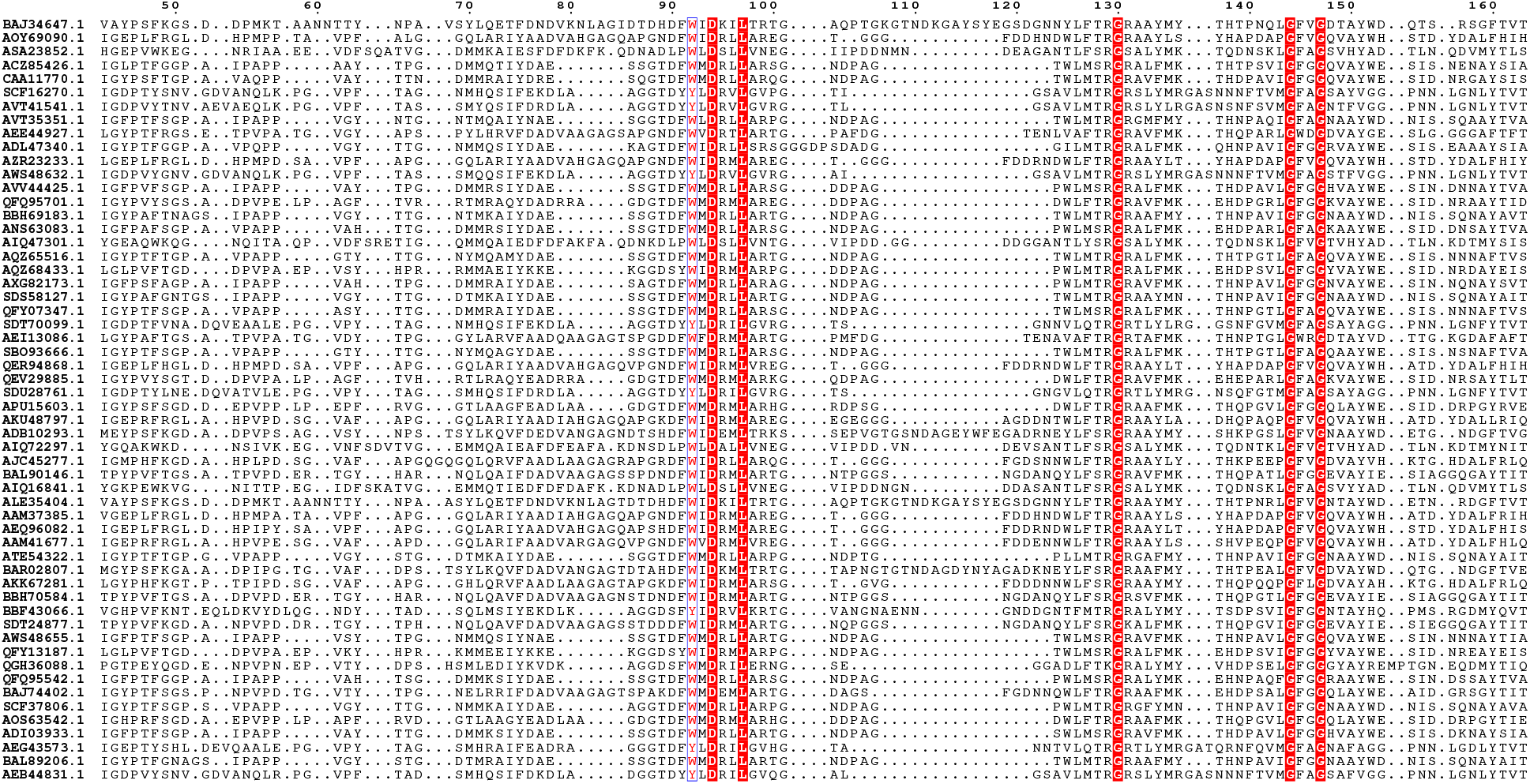
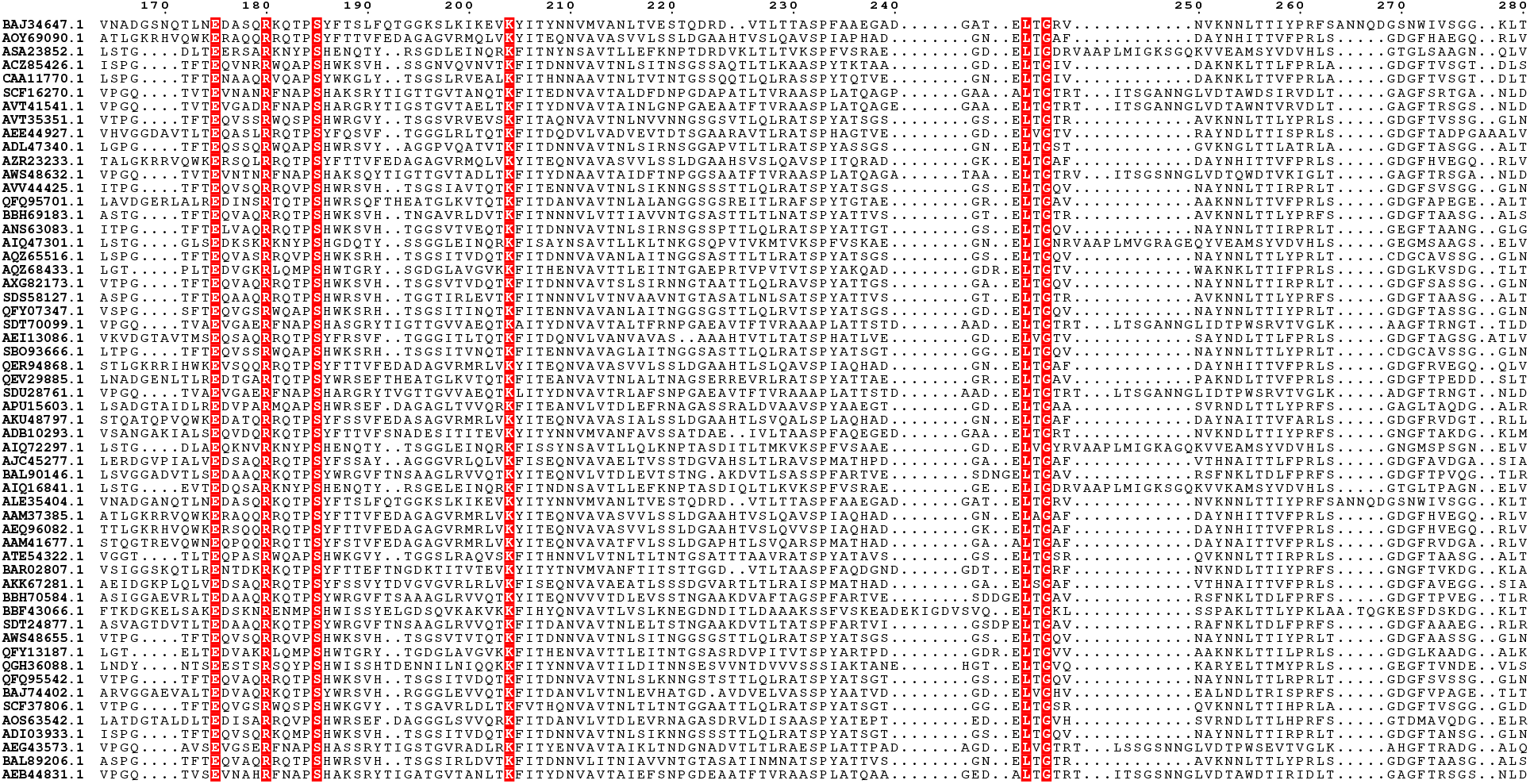
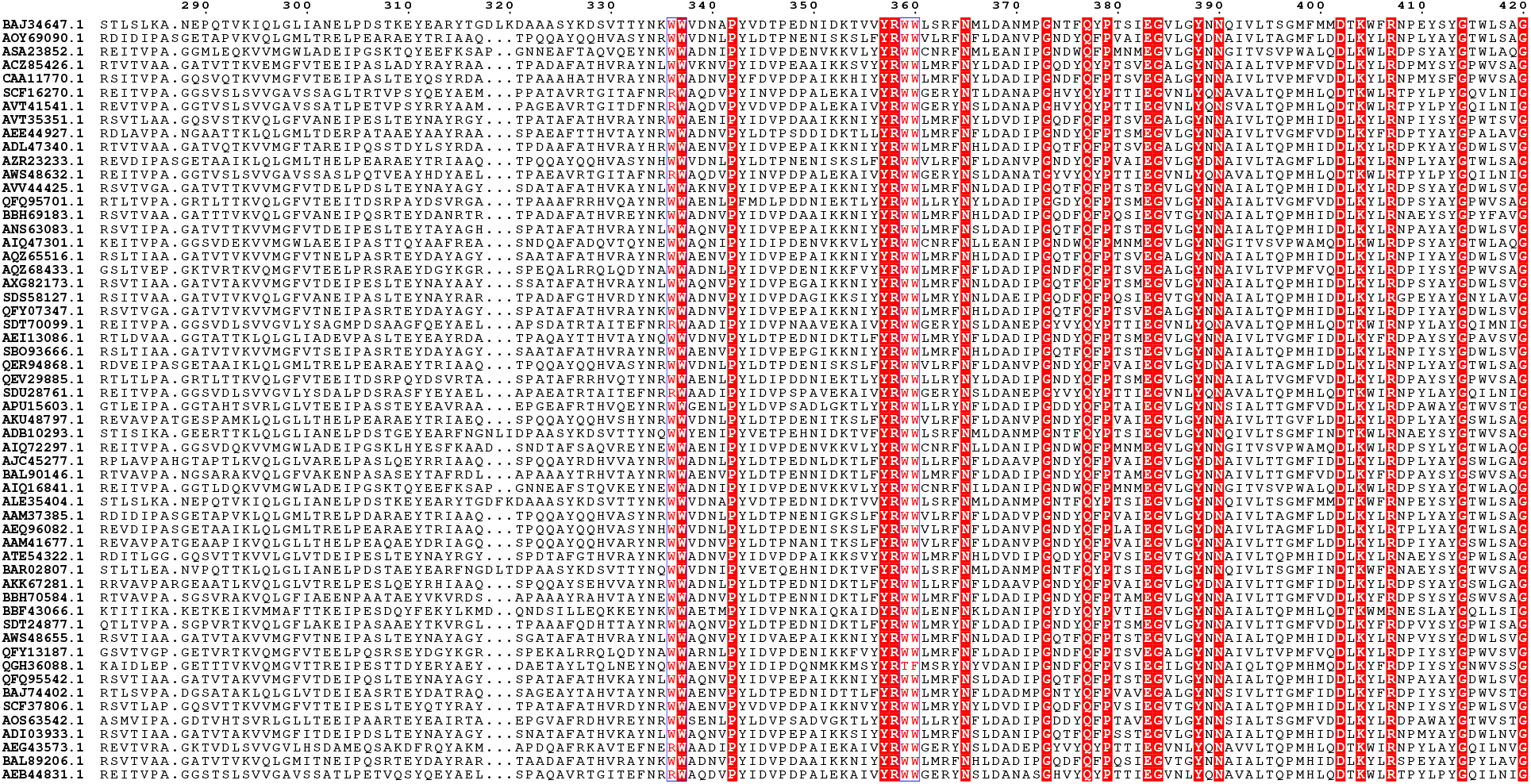
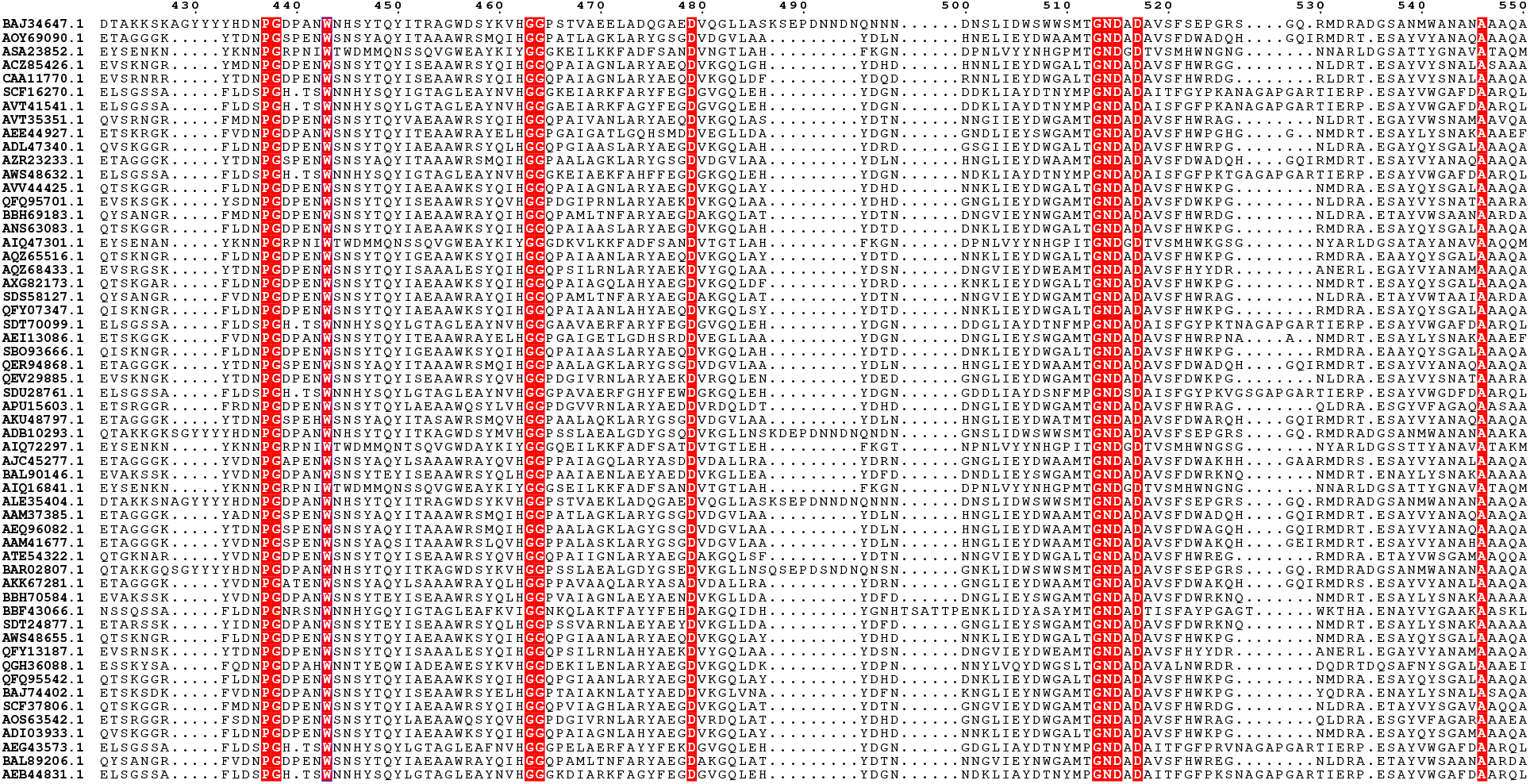
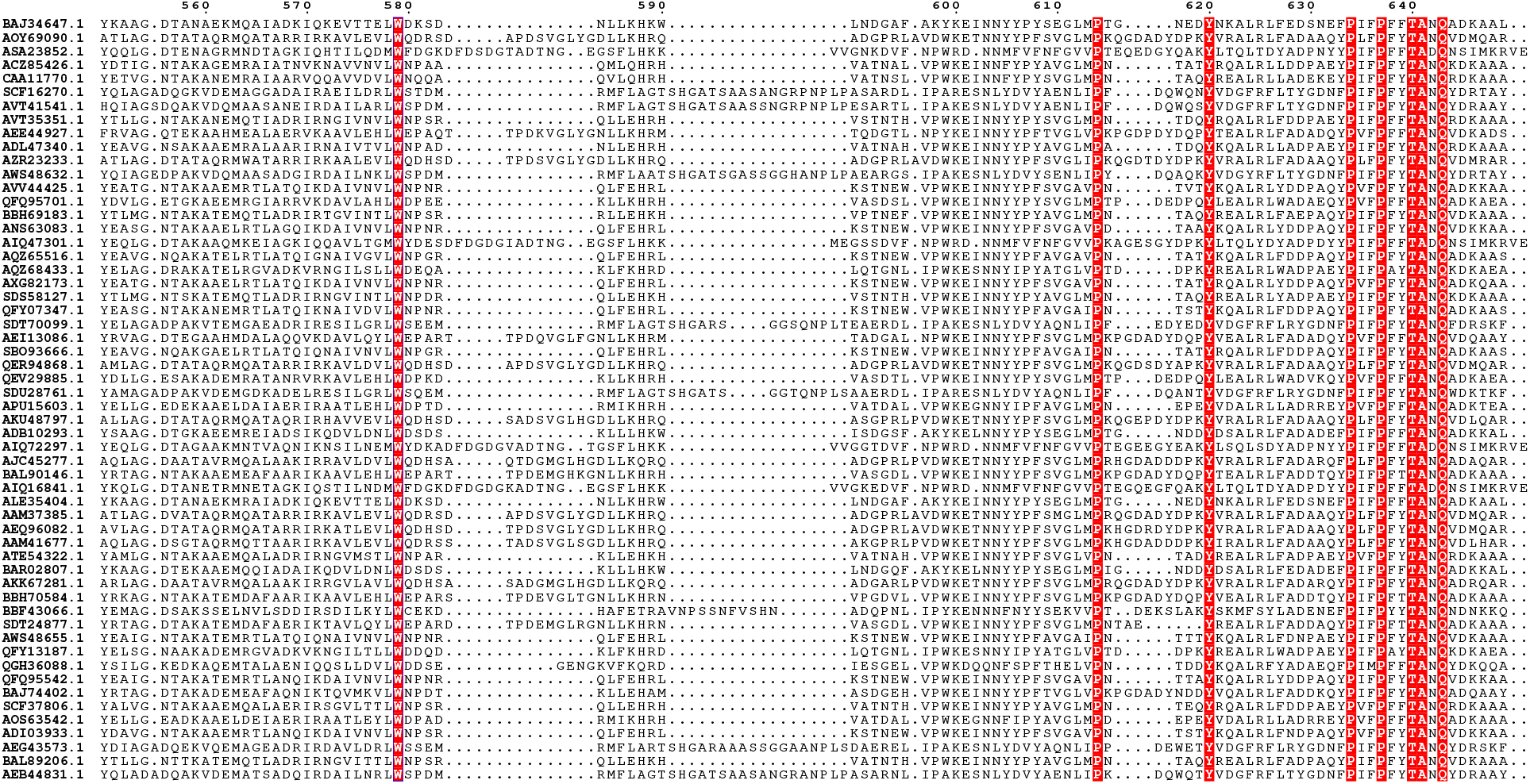
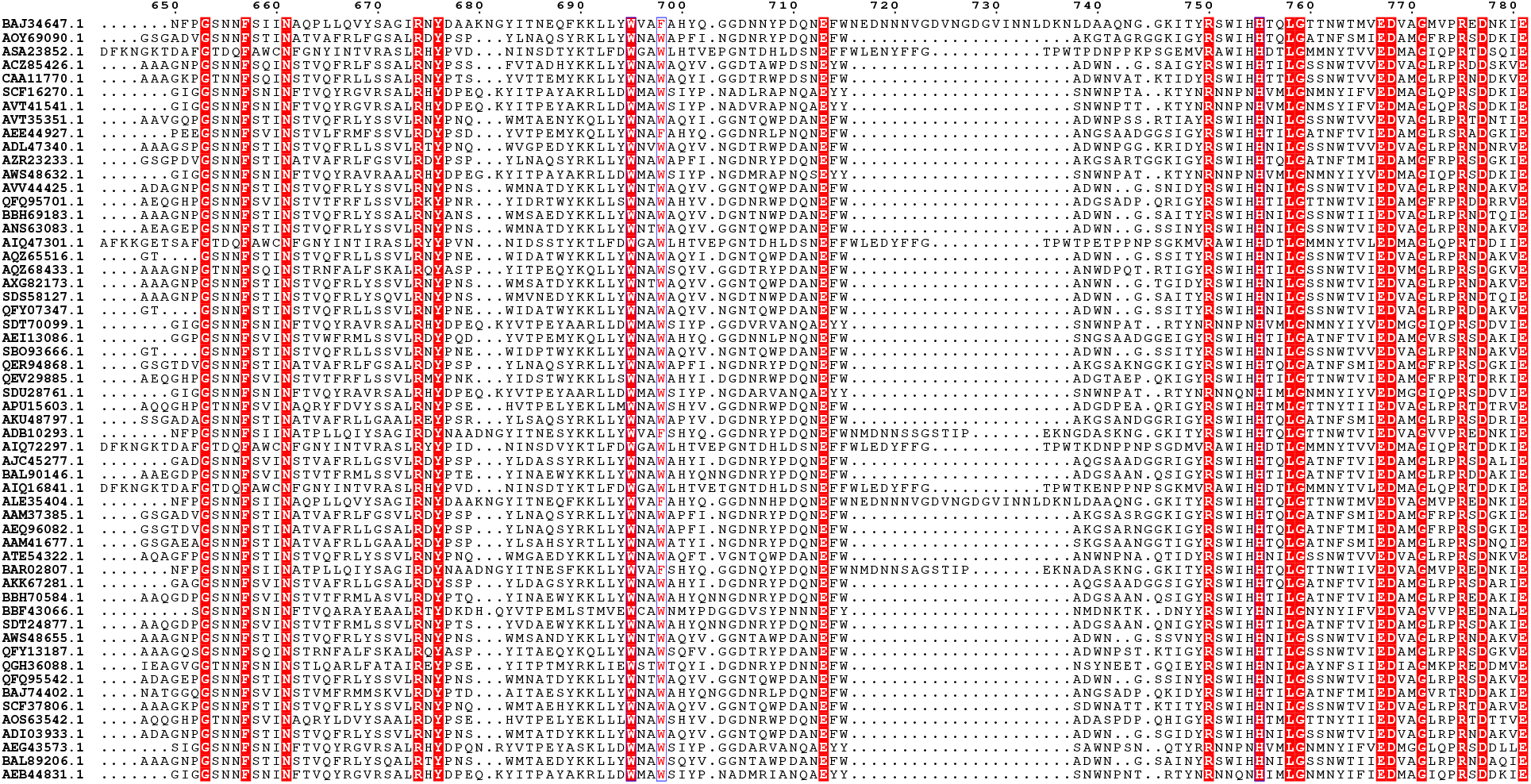
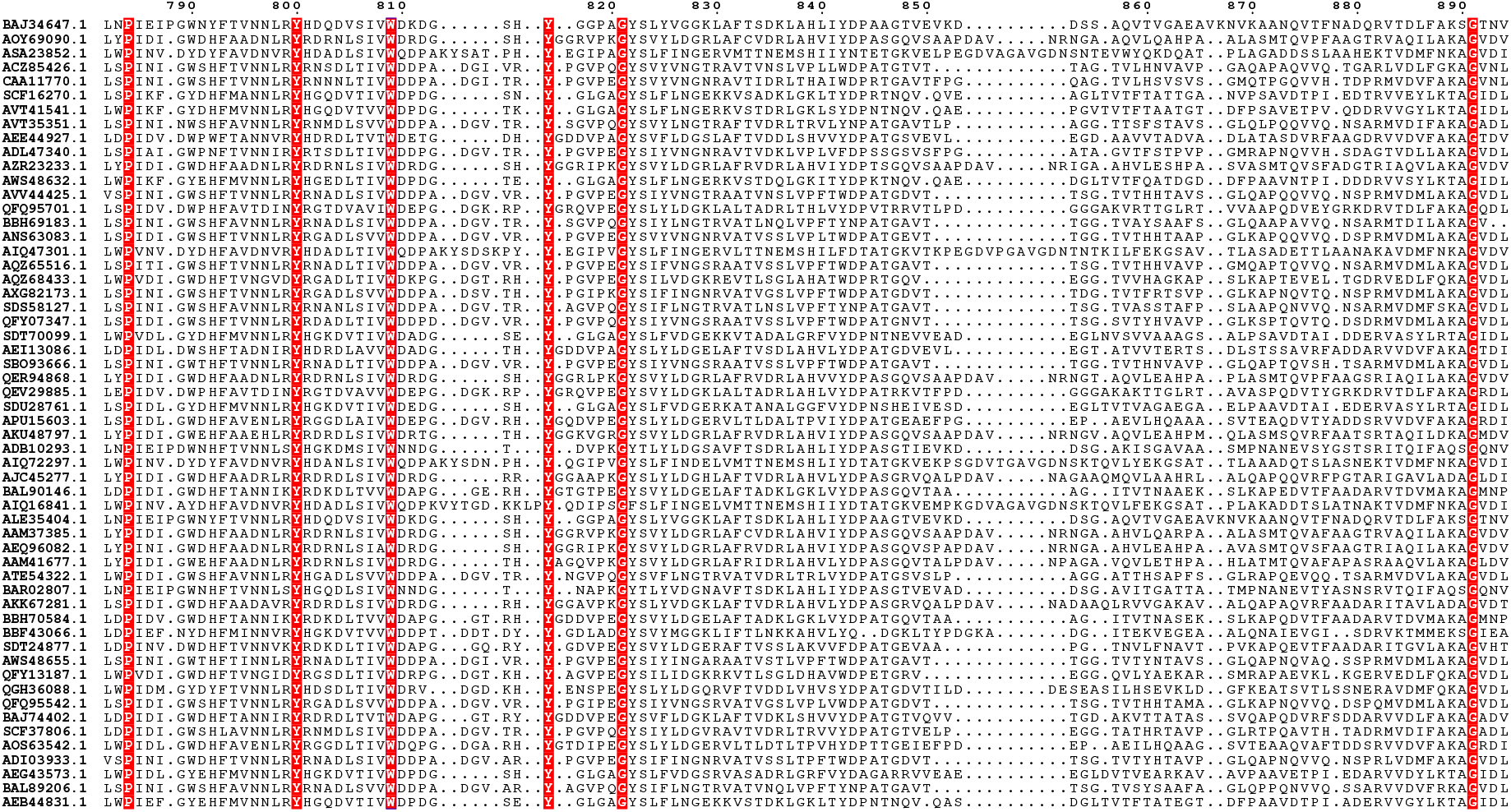
Full amino acid sequence alignment of GH121 entries. From 272 entries of GH121, 56 were selected. HypBA2 is shown at the top (BAJ34647.1).

## References

1. Shallom D, Shoham Y. Microbial hemicellulases. Current Opinion in Microbiology. 2003. pp. 219–228. doi:10.1016/S1369-5274(03)00056-0

2. Garron ML, Henrissat B. The continuing expansion of CAZymes and their families. Current Opinion in Chemical Biology. 2019. pp. 82–87. doi:10.1016/j.cbpa.2019.08.004

3. Kieliszewski MJ, Lamport DTA. Extensin: repetitive motifs, functional sites, post-translational codes, and phylogeny. The Plant Journal. 1994. pp. 157–172. doi:10.1046/j.1365-313X.1994.05020157.x

4. Kieliszewski MJ, Showalter AM, Leykam JF. Potato lectin: a modular protein sharing sequence similarities with the extensin family, the hevein lectin family, and snake venom disintegrins (platelet aggregation inhibitors). Plant J. 1994;5: 849–861. doi:10.1046/j.1365-313X.1994.5060849.x

5. Matsubayashi Y. Posttranslationally modified small-peptide signals in plants. Annu Rev Plant Biol. 2014;65: 385–413. doi:10.1146/annurev-arplant-050312-120122

6. Fujita K, Sakamoto S, Ono Y, Wakao M, Suda Y, Kitahara K, et al. Molecular cloning and characterization of a β-L-arabinobiosidase in *Bifidobacterium longum* that belongs to a novel glycoside hydrolase family. J Biol Chem. 2011;286: 5143–5150. doi:10.1074/jbc.M110.190512

7. Fujita K, Takashi Y, Obuchi E, Kitahara K, Suganuma T. Characterization of a novel β-L-arabinofuranosidase in *Bifidobacterium longum*: Functional elucidation of a DUF1680 protein family member. J Biol Chem. 2014;289: 5240–5249. doi:10.1074/jbc.M113.528711

8. Ndeh D, Rogowski A, Cartmell A, Luis AS, Baslé A, Gray J, et al. Complex pectin metabolism by gut bacteria reveals novel catalytic functions. Nature. 2017;544: 65–70. doi: 10.1038/nature21725

9. Luis AS, Briggs J, Zhang X, Farnell B, Ndeh D, Labourel A, et al. Dietary pectic glycans are degraded by coordinated enzyme pathways in human colonic *Bacteroides*. Nat Microbiol. 2018;3: 210–219. doi:10.1038/s41564-017-0079-1

10. Ito T, Saikawa K, Kim S, Fujita K, Ishiwata A, Kaeothip S, et al. Crystal structure of glycoside hydrolase family 127 β-L-arabinofuranosidase from *Bifidobacterium longum*. Biochem Biophys Res Commun. 2014;447: 32–37. doi:10.1016/j.bbrc.2014.03.096

11. Sakanaka M, Gotoh A, Yoshida K, Odamaki T, Koguchi H, Xiao JZ, et al. Varied pathways of infant gut-associated *Bifidobacterium* to assimilate human milk oligosaccharides: Prevalence of the gene set and its correlation with bifidobacteria-rich microbiota formation. Nutrients. 2020. doi:10.3390/nu12010071

12. Lamport DTA, Miller DH. Hydroxyproline arabinosides in the plant kingdom. Plant Physiol. 1971;48: 454–456. doi:10.1104/pp.48.4.454

13. Akiyama Y, Mori M, Kato K. 13C-NMR analysis of hydroxyproline arabinosides from *Nicotiana tabacum*. Agric Biol Chem. 1980;44: 2487–2489. doi:10.1271/bbb1961.44.2487

14. Ashford D, Desai NN, Allen AK, Neuberger A, O’Neill MA, Selvendran RR. Structural studies of the carbohydrate moieties of lectins from potato *(Solanum tuberosum)* tubers and thorn-apple *(Datura stramonium)* seeds. Biochem J. 1982;201: 199–208. doi:10.1042/bj2010199

15. Kabsch W. XDS. Acta Crystallogr Sect D Biol Crystallogr. 2010;66: 125–132. doi:10.1107/S0907444909047337

16. Adams PD, Afonine P V., Bunkóczi G, Chen VB, Davis IW, Echols N, et al. PHENIX: A comprehensive Python-based system for macromolecular structure solution. Acta Crystallogr Sect D Biol Crystallogr. 2010;66: 213–221. doi:10.1107/S0907444909052925

17. Emsley P, Lohkamp B, Scott WG, Cowtan K. Features and development of Coot. Acta Crystallogr Sect D Biol Crystallogr. 2010;66: 486–501. doi:10.1107/S0907444910007493

18. Murshudov GN, Skubák P, Lebedev AA, Pannu NS, Steiner RA, Nicholls RA, et al. REFMAC5 for the refinement of macromolecular crystal structures. Acta Crystallogr Sect D Biol Crystallogr. 2011;67: 355–367. doi:10.1107/S0907444911001314

19. Holm L, Laakso LM. Dali server update. Nucleic Acids Res. 2016;44: W351–W355. doi:10.1093/nar/gkw357

20. Ashkenazy H, Abadi S, Martz E, Chay O, Mayrose I, Pupko T, et al. ConSurf 2016: an improved methodology to estimate and visualize evolutionary conservation in macromolecules. Nucleic Acids Res. 2016;44: W344–W350. doi:10.1093/nar/gkw408

21. Katoh K, Standley DM. MAFFT multiple sequence alignment software version 7: Improvements in performance and usability. Mol Biol Evol. 2013;30: 772–780. doi: 10.1093/molbev/mst010

22. Waterhouse AM, Procter JB, Martin DMA, Clamp M, Barton GJ. Jalview Version 2-A multiple sequence alignment editor and analysis workbench. Bioinformatics. 2009;25: 1189–1191. doi:10.1093/bioinformatics/btp033

23. Stamatakis A. RAxML version 8: A tool for phylogenetic analysis and post-analysis of large phylogenies. Bioinformatics. 2014;30: 1312–1313. doi: 10.1093/bioinformatics/btu033

24. Letunic I, Bork P. Interactive tree of life (iTOL) v3: an online tool for the display and annotation of phylogenetic and other trees. Nucleic Acids Res. 2016;44: W242–W245. doi: 10.1093/nar/gkw290

25. Dawson NL, Lewis TE, Das S, Lees JG, Lee D, Ashford P, et al. CATH: An expanded resource to predict protein function through structure and sequence. Nucleic Acids Res. 2017;45: D289–D295. doi:10.1093/nar/gkw1098

26. Kurakata Y, Uechi A, Yoshida H, Kamitori S, Sakano Y, Nishikawa A, et al. Structural insights into the substrate specificity and function of </i>Escherichia coli</i> K12 YgjK, a glucosidase belonging to the glycoside hydrolase family 63. J Mol Biol. 2008;381: 116–128. doi:10.1016/j.jmb.2008.05.061

27. Nakamura M, Yasukawa Y, Furusawa A, Fuchiwaki T, Honda T, Okamura Y, et al. Functional characterization of unique enzymes in *Xanthomonas euvesicatoria* related to degradation of arabinofurano-oligosaccharides on hydroxyproline-rich glycoproteins. PLoS One. 2018;13: e0201982. doi:10.1371/journal.pone.0201982

28. Miyazaki T, Nishikawa A, Tonozuka T. Crystal structure of the enzyme-product complex reveals sugar ring distortion during catalysis by family 63 inverting α-glycosidase. J Struct Biol. 2016;196: 479–486. doi:10.1016/j.jsb.2016.09.015

29. Hidaka M, Honda Y, Kitaoka M, Nirasawa S, Hayashi K, Wakagi T, et al. Chitobiose phosphorylase from Vibrio proteolyticus, a member of glycosyl transferase family 36, has a clan GH-L-like (alpha/alpha)(6) barrel fold. Structure. 2004;12: 937–47. doi:10.1016/j.str.2004.03.027

30. Fujimoto Z, Jackson A, Michikawa M, Maehara T, Momma M, Henrissat B, et al. The structure of a *Streptomyces avermitilis* α-L-Rhamnosidase reveals a novel carbohydrate-binding module CBM67 within the six-domain arrangement. J Biol Chem. 2013;288: 12376–12385. doi:10.1074/jbc.M113.460097

31. Gibson RP, Gloster TM, Roberts S, Warren RAJ, Storch De Gracia I, García Á, et al. Molecular basis for trehalase inhibition revealed by the structure of trehalase in complex with potent inhibitors. Angew Chemie - Int Ed. 2007;46: 4115–4119. doi:10.1002/anie.200604825

32. Hidaka M, Kitaoka M, Hayashi K, Wakagi T, Shoun H, Fushinobu S. Structural dissection of the reaction mechanism of cellobiose phosphorylase. Biochem J. 2006;398: 37–43. doi:10.1042/BJ20060274

33. Nerinckx W, Desmet T, Piens K, Claeyssens M. An elaboration on the syn-anti proton donor concept of glycoside hydrolases: Electrostatic stabilisation of the transition state as a general strategy. FEBS Lett. 2005;579: 302–312. doi:10.1016/j.febslet.2004.12.021

34. The CAZypedia Consortium. Ten years of CAZypedia: a living encyclopedia of carbohydrate-active enzymes. Glycobiology. 2018;28: 3–8. doi:10.1093/glycob/cwx089

35. Hasegawa K, Kubota M, Matsuura Y. Roles of catalytic residues in α-amylases as evidenced by the structures of the product-complexed mutants of a maltotetraose-forming amylase. Protein Eng. 1999;12: 819–824. doi:10.1093/protein/12.10.819

36. Matsuura Y. A possible mechanism of catalysis involving three essential residues in the enzymes of α-amylase family. Biologia - Section Cellular and Molecular Biology. 2002. pp. 21–27.

37. Brzozowski AM, Davies GJ. Structure of the *Aspergillus oryzae* α-amylase complexed with the inhibitor acarbose at 2.0 Å resolution. Biochemistry. 1997;36: 10837–10845. doi:10.1021/bi970539i

38. Sakurama H, Fushinobu S, Hidaka M, Yoshida E, Honda Y, Ashida H, et al. 1,3-1,4-α-L-Fucosynthase that specifically introduces Lewis a/x antigens into type-1/2 chains. J Biol Chem. 2012. doi:10.1074/jbc.M111.333781

39. Ito T, Katayama T, Hattie M, Sakurama H, Wada J, Suzuki R, et al. Crystal structures of a glycoside hydrolase family 20 lacto-*A*-biosidase from *Bifidobacterium bifidum*. J Biol Chem. 2013;288: 11795–11806. doi:10.1074/jbc.M112.420109

40. Yamada C, Gotoh A, Sakanaka M, Hattie M, Stubbs KA, Katayama-Ikegami A, et al. Molecular insight into evolution of symbiosis between breast-fed infants and a member of the human gut microbiome *Bifidobacterium longum*. Cell Chem Biol. 2017;24: 515–524.e5. doi:10.1016/j.chembiol.2017.03.012

41. Suzuki R, Katayama T, Kitaoka M, Kumagai H, Wakagi T, Shoun H, et al. Crystallographic and mutational analyses of substrate recognition of endo-alpha-N-acetylgalactosaminidase from *Bifidobacterium longum*. J Biochem. 2009;146: 389–98. doi:10.1093/jb/mvp086

42. Tian W, Chen C, Lei X, Zhao J, Liang J. CASTp 3.0: Computed atlas of surface topography of proteins. Nucleic Acids Res. 2018;46: W363–W367. doi: 10.1093/nar/gky473

43. El-Gebali S, Mistry J, Bateman A, Eddy SR, Luciani A, Potter SC, et al. The Pfam protein families database in 2019. Nucleic Acids Res. 2019;47: D427–D432. doi: 10.1093/nar/gky995

44. Miyazaki T, Ichikawa M, Iino H, Nishikawa A, Tonozuka T. Crystal structure and substrate-binding mode of GH63 mannosylglycerate hydrolase from *Thermus thermophilus* HB8. J Struct Biol. 2015. doi:10.1016/j.jsb.2015.02.006

45. Pachl P, Škerlová J, Šimčíková D, Kotik M, Křenková A, Mader P, et al. Crystal structure of native α-L-rhamnosidase from *Aspergillus terreus*. Acta Crystallogr Sect D Struct Biol. 2018;74: 1078–1084. doi:10.1107/S2059798318013049

46. Cardona F, Parmeggiani C, Faggi E, Bonaccini C, Gratteri P, Sim L, et al. Total syntheses of casuarine and its 6-*O*-α-glucoside: Complementary inhibition towards glycoside hydrolases of the GH31 and GH37 families. Chem - A Eur J. 2009;15: 1627–1636. doi:10.1002/chem.200801578

47. Cereija TB, Alarico S, Lourenço EC, Manso JA, Ventura MR, Empadinhas N, et al. The structural characterization of a glucosylglycerate hydrolase provides insights into the molecular mechanism of mycobacterial recovery from nitrogen starvation. IUCrJ. 2019;6: 572–585. doi:10.1107/S2052252519005372

48. Nam YW, Nihira T, Arakawa T, Saito Y, Kitaoka M, Nakai H, et al. Crystal structure and substrate recognition of cellobionic acid phosphorylase, which plays a key role in oxidative cellulose degradation by microbes. J Biol Chem. 2015. doi:10.1074/jbc.M115.664664

49. Bianchetti CM, Elsen NL, Fox BG, Phillips GN. Structure of cellobiose phosphorylase from Clostridium thermocellum in complex with phosphate. Acta Crystallogr Sect F Struct Biol Cryst Commun. 2011. doi:10.1107/S1744309111032660

50. Barker MK, Rose DR. Specificity of processing α-glucosidase I is guided by the substrate conformation: Crystallographic and in silico studies. J Biol Chem. 2013;288: 13563–13574. doi:10.1074/jbc.M113.460436

